# In vitro study of chemical compositions and antibacterial activity of Lorestan endemic plant, *Pistacia atlantica* leaf hydroalcoholic extract against cariogenic bacteria

**DOI:** 10.1101/2024.09.29.615670

**Authors:** Mahnaz Rahimi, Sara Mohammadi Sepah Vand, Azadeh Mohammadi Sepahvand

## Abstract

**Background and objectives:** The use of medicinal plants has been considered as an adjunctive treatment to prevent and control dental caries, alongside the mechanical removal of microbial agents. The aim of this study was to evaluate the chemical compositions and antibacterial activity of *Pistacia atlantica* leaf hydroalcoholic extract against standard strains of *Streptococcus mutans (SM)* and *Lactobacillus rhamnosus (LR)*.

**Materials and Methods:** The hydroalcoholic extract of *Pistacia atlantica* leaves was prepared using the maceration method, and the chemical compositions were identified by a GC-MS apparatus. The antibacterial activity was assessed on the standard strains of *SM* and *LR* by measuring the mean diameter of the inhibition zone at different concentrations using the agar well diffusion method, and determining the minimum inhibitory concentration (MIC) and minimum bactericidal concentration (MBC) using the broth microdilution method. Chlorhexidine (CHX) mouthwash at a concentration of 0.2% was used as the positive control group. The tests were repeated three times, and the results were analyzed using ANOVA, Tukey’s Honestly Significant Difference (HSD) test, and T-test with SPSS. A significance level of P< 0.05 was considered.

**Results:** The MIC of the *Pistacia atlantica* leaf hydroalcoholic extract for *SM, LR*, and CHX was 12.5, 25, and 6.25 mg/ml, respectively, and the MBC for *SM, LR*, and CHX was 25, 25, and 6.25 mg/ml, respectively. In the agar well diffusion method, the mean diameter of the inhibition zone at the concentration of 200 mg/ml for *SM, LR*, and CHX was 23.33±6.50, 19.0±1.0, and 17.33±0.57 mm, respectively. The major chemical compositions were *α* -pinene, *α* -bisabolene, 3-hexenol, pulegone, and *β*-pinene, respectively.

**Conclusion:** The leaf hydroalcoholic extract of *Pistacia atlantica* demonstrated antibacterial activity against *SM* and *LR*.

## Introduction

Oral infections are one of the most common human infections (Tahmourespour and Kermanshahi 2011). Among these, dental caries and gingival diseases are the most prevalent types, which are caused by the accumulation of microorganisms and formation of microbial plaque (Zhang et al. 2018). According to RC. Shields, almost 2.4 billion adults and 621 million children had experienced untreated dental caries by 2010 (Shields et al. 2018). Dental caries results from the extended interaction of cariogenic microorganisms, particularly *Streptococcus mutans* and *Lactobacillus rhamnosus*, with a diet abundant in fermentable carbohydrates, along with various host factors such as salivary secretion levels and buffer capacity (Cagetti et al. 2013). *Streptococcus mutans*, a gram-positive coccus-shaped acidogenic bacterium, is among the most important microorganisms present in dental plaque, playing a significant role in the onset of dental caries, while *Lactobacillus rhamnosus*, another gram-positive bacterium, typically residing in the digestive system, plays an essential role in the progression of dental caries (Ghasemi et al. 2016; Widyarman, Enrita Dian, and Fibryanto 2022; Loesche 1996). A common approach for reducing the number of cariogenic bacteria is by mechanically removing dental caries (Avinash et al. 2012). However, it is important to acknowledge that tooth restoration can be costly. Consequently, the ability to afford these expenses is likely the most significant obstacle to receiving restorative dental treatments (Nahvi et al. 2017). Some of the most common methods evaluated in recent research to mitigate imposed costs include the use of antibacterial materials, antibiotics, fluoride therapy, vaccination, and medicinal plants with antibacterial activity against cariogenic bacteria (F. Ahrari et al. 2016; Khoroushi, Khorasgani, and Aliasghari 2017; Caries 2017). The utilization of medicinal plants in traditional medicine remains popular, particularly in developing countries (Che et al. 2017). Several studies have indicated that certain plants containing compounds such as flavonoids and other polyphenols, terpenes, and alkaloids may exhibit antimicrobial activities against oral microorganisms (Khoroushi, Khorasgani, and Aliasghari 2017; Cowan 1999; Othman, Sleiman, and Abdel-Massih 2019; Álvarez-Martínez et al. 2021). This form of treatment has deep cultural roots in regions such as Iran, with a longstanding history of traditional medicine practices (Buso et al. 2020). Delfan et al. reported a list of 14 medicinal plants collected from eight cities in Lorestan province, extensively used by locals to alleviate tooth pain, among which *Pistacia atlantica* stands out as available to be collected in all seasons (B. Delfan et al. 2014). This native plant, belonging to the Anacardiaceae family, thrives in Iran’s Zagros region and western areas (Ghasemi et al. 2016). In contemporary times, there is a notable resurgence in public preference for using medicinal plants in oral hygiene products, particularly in developed countries (Karunamoorthi et al. 2013; Tapsoba and Deschamps 2006). As a result, many pharmaceutical and hygienic factories have begun incorporating extracts of medicinal plants into the production of toothpastes and mouthwashes (Rajendiran et al. 2021; Suresh et al. 2021). *Pistacia atlantica*, comprises approximately 70 genera and over 600 species (Bozorgi et al. 2013). Scientific evidence has demonstrated that different parts of *Pistacia species* exhibit medicinal effects (Ghasemi et al. 2016). Traditionally, the aerial parts of the plant have been used as a stimulant for their diuretic properties in treating hypertension, as well as other conditions such as coughs, sore throat, eczema, and stomachaches (Omidi and Sharifi 2017). The green-colored, bitter-tasting gum of *Pistacia atlantica* is utilized to heal wounds, alleviate toothaches and kidney pain, and treat infections (Soleiman-Beigi and Arzehgar 2013). Among its effects, anti-oxidant, anti-viral, anti-tumoral, and anti-inflammatory properties are noteworthy (Benmohamed et al. 2023; Najjari et al. 2022). Studies have revealed that the gum of *Pistacia* species exerts antimicrobial effects against Staphylococcus aureus, Salmonella enteritidis, Bacillus cereus, Escherichia coli, Helicobacter pylori, mildew, and yeast (Kermanshah et al. 2011). Moreover, the leaves of this plant possess significant anti-viral, antimicrobial, and antifungal activities. Various studies have demonstrated these activities against Staphylococcus aureus, Listeria monocytogenes, Pseudomonas aeruginosa, Escherichia coli, Salmonella typhi, Bacillus cereus, Candida albicans, and others (Benhammou, Bekkara, and Panovska 2008; Barzegar, Hojjati, and Panahi 2017). Chemical compositions including *α* -pinene, limonene, linalool, and terpinene have been identified as responsible for the mentioned effects (Soković and Van Griensven 2006; Mahmoudvand et al. 2016). This study aimed to assess the chemical composition and in vitro antibacterial activity of the hydroalcoholic extract from the leaves of *Pistacia atlantica*, an endemic plant in Lorestan, against cariogenic bacteria. The antimicrobial effects of this plant against cariogenic bacteria could serve as a cost-effective strategy for the prevention and control of dental caries.

## Methods and Materials

### Plant collection and extraction

First, leaves of *Pistacia atlantica* were collected from the mountains of Lorestan province in October 2022. After identification and scientific verification, the samples were dried in the shade, away from direct sunlight, and then powdered. The powder was preserved in the refrigerator for future applications. Extraction was performed using the maceration method. To prepare the hydroalcoholic extract, 15 grams of leaf powder, weighed by a digital weighing machine (Sartorius TE64 Germany), were mixed with a specific amount of ethanol 70% as a solvent. Subsequently, the resulting solution was kept in an incubator at 60 °C for 48 hours and then centrifuged at 10,000 rpm for 10 minutes. After repeating this procedure twice, the filtered solution was concentrated using a rotary evaporator (Heidolph, Germany). Finally, the concentrated solution was transferred to an oven and dried at 60 °C before being stored in the refrigerator for further experiments.

### Bacteria preparation

Standard strains of *Streptococcus mutans* (PTCC:1683) and *Lactobacillus rhamnosus* (PTCC:1637) were acquired from the Iranian Research Organization for Science and Technology in Tehran, Iran. The samples were cultured in nutrient broth medium for 24 hours at 30-35 °C. Subsequently, the bacteria were cultured on blood agar culture for *Streptococcus mutans* and MRS agar for *Lactobacillus rhamnosus* following the provided instructions.

### Measuring the inhibition zone

A suspension of each bacteria mixed with saline was prepared to achieve a turbidity equivalent to 0.5 McFarland standards (1.5×108 CFU ml-1). Then, suspensions of *Streptococcus mutans* and *Lactobacillus rhamnosus* were spread on the surfaces of blood agar and MRS agar plates, respectively, using sterile cotton swabs. The agar media were then punched with 6 mm diameter wells. Next, 100 *µ*l of the extract with concentrations of 10, 50, 100, and 200 mg/ml was injected into the wells. Dimethyl sulfoxide (DMSO) disks served as the negative control group, while CHX mouthwash 0.2% was used as the positive control group. Finally, both agar plates were incubated at 37 °C for 24 hours. The inhibition zones around the wells were measured in millimeters.

### Determination of minimum inhibitory concentration (MIC) and minimum bactericidal concentration (MBC)

To evaluate the antibacterial activity of the hydroalcoholic extract from *Pistacia atlantica* leaf against cariogenic bacteria, the Broth microdilution method was employed. Initially, 100 *µ*l of brain heart infusion (BHI) was dispensed into each well of a 96-cell culture plate. Then, 100 *µ*l of the extract was added to the first well and thoroughly mixed. Subsequently, 100 *µ*l of the content from the first well was transferred to the second well, and this process was repeated sequentially for each well. Eventually, 100 *µ*l of the content from the last well was discarded. Different dilutions were prepared from the extract. BHI environments with DMSO and CHX served as the negative and positive control groups, respectively. The samples were then incubated in an incubator for 18-24 hours at 37 °C. The minimum concentration of the extract that resulted in no visible bacterial growth (turbidity) was considered the MIC. To determine the MBC, samples showing no turbidity were inoculated onto MRS agar for *Lactobacillus rhamnosus* and blood agar for *Streptococcus mutans*. The agar plates were then incubated at 37 °C for 24 hours. The lowest concentration of the extract capable of killing at least 99.9% of the bacteria was considered the MBC. Each experiment was repeated three times.

### Chromatographic analysis

For GC-MS analysis, a Shimadzu model GC-17A (Kyoto, Japan) gas chromatograph coupled to a Shimadzu Quadruple-MS model QP5050 mass spectrometer was utilized. Compounds were separated on a 30 m × 0.22 mm i.d. fused-silica capillary column coated with a 0.25 µm film of BP-5 (Shimadzu) using a split/splitless injector with a 1 mm internal diameter glass liner. Ultra-pure helium was employed as the carrier gas with an ionization voltage of 70 eV. The injector and interface temperatures were maintained at 280 °C and 260 °C, respectively. The mass ranged from 35 to 450 amu. The oven temperature program was the same as mentioned earlier for the GC. The constituents of the extract were identified by calculating their retention indices under temperature-programmed conditions for n-alkanes (C8–C20) and the extract on a DB-5 column under the same chromatographic conditions. Identification of genotype compounds was conducted by comparing their mass spectra with those of the internal reference mass spectra library (NIST08 and Wiley 9.0)

### Statistical analysis

One-way ANOVA, Tukey’s HSD, and T-test were used in search of statistically significant differences. Data analysis was performed using the SPSS 22.0 software (SPSS Inc., Chicago, IL, USA). The significance level was considered for a P< 0.05.

## Results

Table 1 indicates the results obtained from GC-MS analysis of the hydroalcoholic extract from *Pistacia atlantica* leaves. (Components comprising less than 4% of the total identified compounds were not mentioned). The main components of the extract were *β* -pinene, *β* - bisabolene, 3 -hexenol, pulegone, *β* pinene, bornyl acetate and spathulenol.

**Table 1.** Chemical components and their percentages in P.atlantica leaf hydroalcoholic extract; t_R: Retention time(min), RI : Retention index.

| $t_R$ | RI | Component | Area(%) |
| --- | --- | --- | --- |
| 4.858 | 905.524 | α-pinene | 27.09 |
| 22.4 | 1502.544 | α-bisabolene | 18.71 |
| 3.625 | 788.095 | 3-hexenol | 6.93 |
| 12.8 | 1247.864 | pulegone | 5.85 |
| 5.725 | 961.667 | β-pinene | 5.82 |
| 14.192 | 1288.866 | bornyl acetate | 5.43 |
| 25.767 | 1588.219 | spathulenol | 4.42 |

Retention time (*t*_*R*_) measures a solute’s passage through a column, from injection to detection. *t*_*R*_ is not entirely exclusive, as multiple substances can share the same *t*_*R*_ due to operational factors like carrier gas type and flow(Bizzo et al. 2023). To address this limitation in compound identification, Retention index (RI) values are used. These values, introduced by Kováts, provide a standardized approach alongside mass spectrometry, overcoming these shortcomings(Tarjan et al. 1989).

Table 2 indicates the mean diameter of the inhibition zone of the hydroalcoholic extract from *Pistacia atlantica* leaves at different concentrations against *Streptococcus mutans, Lactobacillus rhamnosus*, and the control group (in millimeters).

**Table 2.**
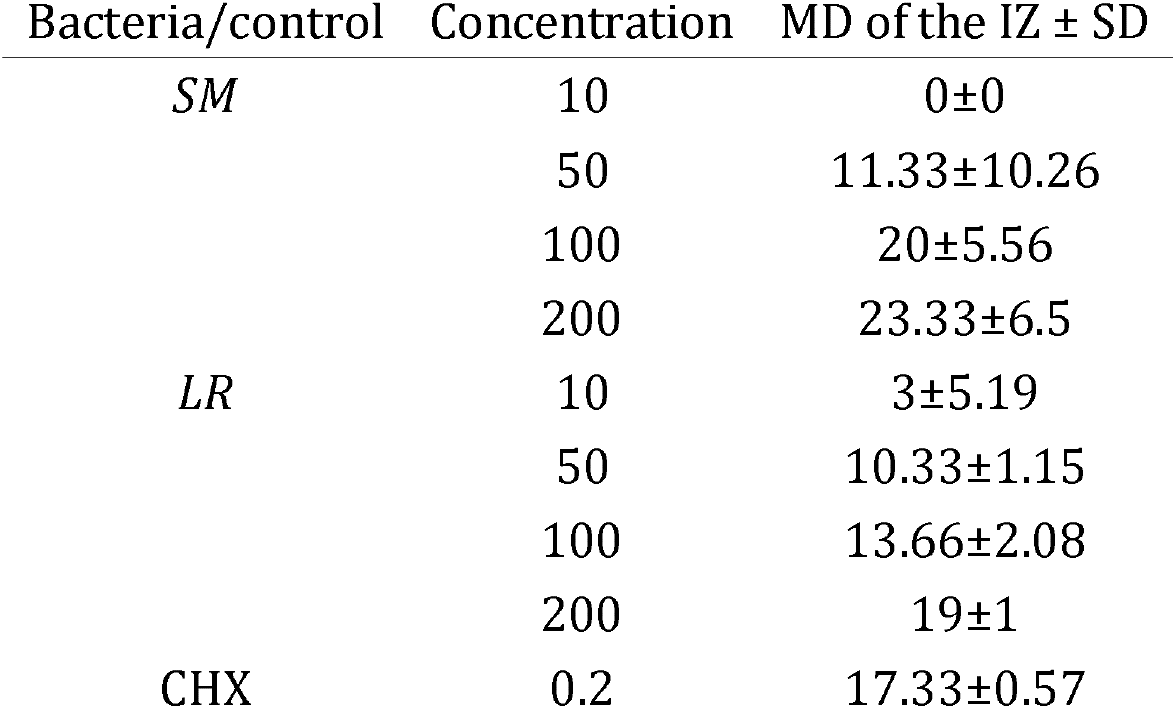
The mean diameter (MD) and standard deviation (SD) of the inhibition zone (IZ) in millimeters at different concentrations in mg/ml of the hydroalcoholic extract from Pistacia atlantica leaf against Streptococcus mutans, Lactobacillus rhamnosus, and Chlorhexidine.

| Bacteria/control | Concentration | MD of the IZ ± SD |
| --- | --- | --- |
| SM | 10 | 0±0 |
| | 50 | 11.33±10.26 |
| | 100 | 20±5.56 |
| | 200 | 23.33±6.5 |
| LR | 10 | 3±5.19 |
| | 50 | 10.33±1.15 |
| | 100 | 13.66±2.08 |
| | 200 | 19±1 |
| CHX | 0.2 | 17.33±0.57 |

The MIC and MBC of *Pistacia atlantica* leaf extract against *Streptococcus mutans, Lactobacillus rhamnosus*, and the control group are reported in Table 3 (in mg/ml).

**Table 3.** The MIC and MBC of the hydroalcoholic extract from Pistacia atlantica leaf against Streptococcus mutans and Lactobacillus rhamnosus in mg/ml.

| Bacteria | MIC | MBC |
| --- | --- | --- |
| *SM* | 12.5 | 25 |
| *LR* | 25 | 25 |
| Positive control | 6.25 | 6.25 |

Figures 1(a) and 1(b) show the mean diameter of the inhibition zone of the hydroalcoholic extract from *Pistacia atlantica* leaves for *Streptococcus mutans* and *Lactobacillus rhamnosus* at concentrations of 100 and 200 mg/ml.

**Fig 1 legend:**
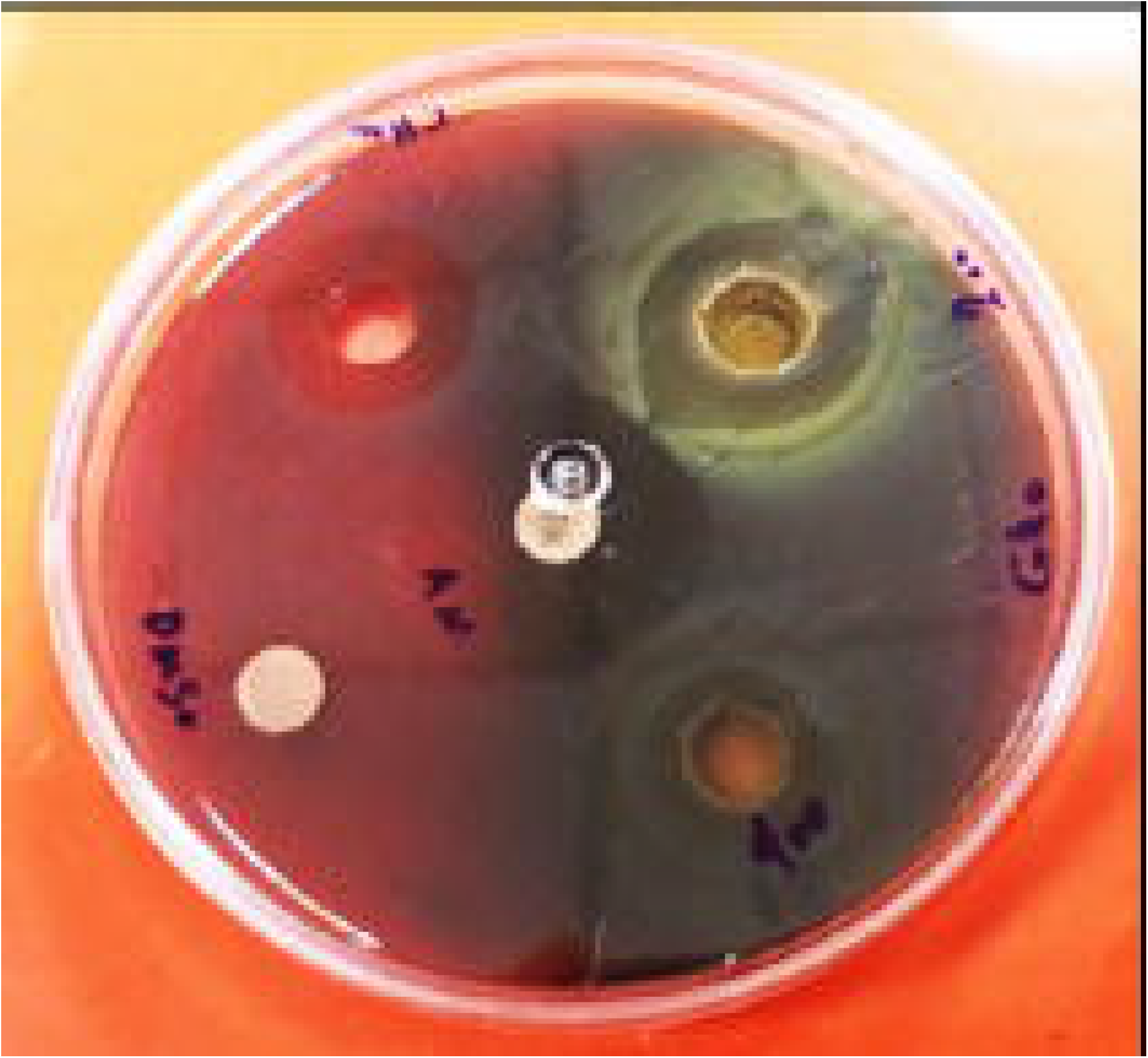
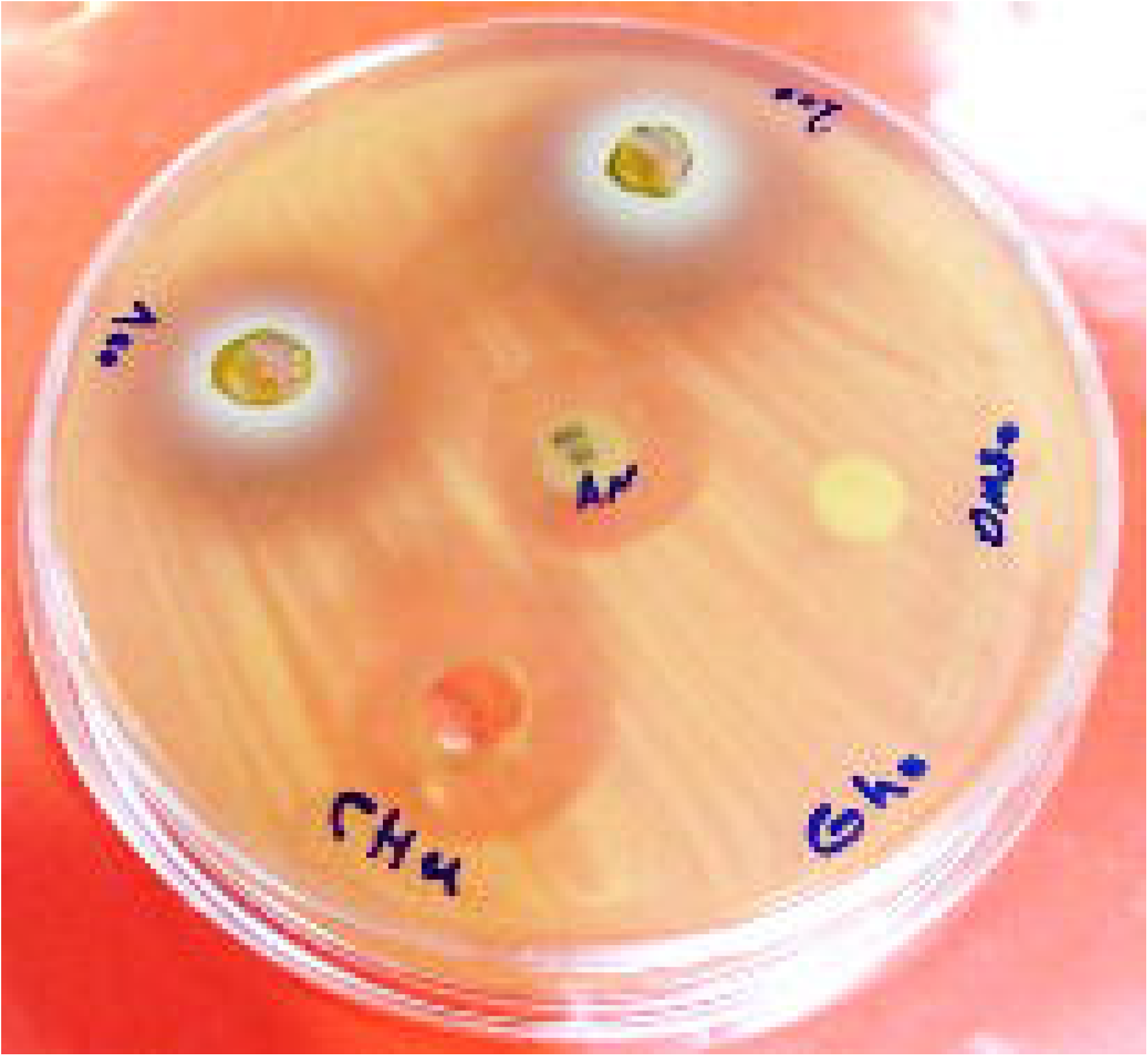
The antibacterial activity of P.atlantica leaf hydroalcoholic extract. The mean diameter of the inhibition zone for (a) *Streptococcus mutans* and (b) *Lactobacillus rhamnosus* at concentrations of 100 and 200 mg/ml in the presence of DMSO and distilled water as the negative control, and CHX mouthwash as the positive control.

The results of ANOVA tests indicated significant differences among groups. To further elucidate these differences, Tukey’s HSD and T-tests were employed.

Regarding the extract’s performance at various concentrations in the presence of *Streptococcus mutans*, no significant differences in the inhibition zone diameter were observed among concentrations of 50, 100, and 200, as well as CHX, except at the 10 mg/ml concentration (P value > 0.05). Notably, the non-growth zones’ diameters at concentrations of 100 and 200 were substantially larger than those observed with CHX. Conversely, when dealing with *Lactobacillus rhamnosus* bacteria, all groups exhibited significant effectiveness compared to the extract at a concentration of 10 mg/ml. However, no significant differences were noted among the extracts at higher concentrations when compared to each other or to CHX.

## Discussion

The adverse effects of certain antibiotics and the widespread presence of multi-drug-resistant bacterial strains pose significant therapeutic challenges. Employing antibiotic-resistant inhibitors derived from medicinal plants is one of the contemporary strategies aimed at mitigating antibiotic resistance (Pratap et al. 2012). Additionally, factors such as the availability of medicinal plants, the better compatibility of specific chemical compounds within these plants with the human immune system, and the patients’ interest towards traditional treatments have contributed to the growing utilization of medicinal plants in addressing health issues(Azmoudeh, Aslanimehr, and Lourizadeh 2017).

Delfan et al. reported that in Lorestan province, many native people use 14 medicinal plant species to relieve their toothache, and *Pistacia atlantica* was among them, notable for being available for collection throughout all seasons(B. Delfan et al. 2014). The authors of this article did not find any in vivo studies on the anti-pain or anti-inflammatory activity of this plant against dental diseases. However, several in vitro studies have been conducted where researchers investigated the antimicrobial effects of various essential oils and extracts (hydroalcoholic, aqueous, etc.) of this plant, collected from different regions, against the cariogenic bacteria. (Roozegar et al. 2016; Tahmourespour, Aminzadeh, and Salehifard 2022; El Zerey-Belaskri, Belyagoubi-Benhammou, and Benhassaini 2022)

Chemical composition: Medicinal plants contain a variety of chemical compounds such as alkaloids, tannins, flavonoids, etc., which possess potential antibacterial properties against cariogenic bacteria through diverse mechanisms. These mechanisms include direct effects on oral bacteria, prevention of bacterial attachment to the tooth surface, and inhibition of glucans and amylase production(Pratap et al. 2012). The antibacterial activity of certain chemical compounds from *Pistacia atlantica* has been reported in previous studies. For instance, the methanolic fruit extracts of Pistacia atlantica, as reported in the study by Mahmoudvand et al., contained *β*-myrcene (41.4%), a-pinene (32.48%), and limonene (4.66%)(Mahmoudvand et al. 2016). Additionally, the essential oil of *Pistacia atlantica* analyzed in the study by Hosseini et al. revealed the presence of *β*-pinene (70%), *β* - copaene (76%), and *β* -terpinolene (86%)(F. Hosseini, Adlgostar, and Sharifnia 2013). The results of the chemical composition analysis in this study revealed that the major components of the extract were a-pinene (27.09%), a-bisabolene (18.71%), 3-hexenol (6.93%), pulegone (5.85%), and *β*-pinene (5.82%). These results demonstrate that a plant species grown in different regions may contain similar or different compounds in varying proportions, ultimately leading to differences in effectiveness. In fact, climatic conditions, growing time and the location of plant harvesting are indeed crucial factors that influence the outcomes of these types of studies. These variables are responsible for significant changes in the composition and, consequently, the effects of plants. Furthermore, the final properties of products may vary depending on factors such as which part of the plant is used, the method of preparation (essential oil or extract), and the type of extract obtained (aqueous, hydroalcoholic, methanolic, etc.). In this study, among the main chemical compounds of *Pistacia atlantica* leaf hydroalcoholic extract, *β* -pinene, pulegone, and *β*-pinene were found to possess antibacterial activity.

The antibacterial activity: The findings of this study indicate that the hydroalcoholic extract from *Pistacia atlantica* leaf exhibits antibacterial effects against *Streptococcus mutans* and Lactobacillus rhamnosus. Several medicinal plants have been evaluated for their antibacterial properties, both within and outside Iran, with reports of effectiveness against *Streptococcus mutans* and *Lactobacillus rhamnosus* using the diffusion method(Daneshkazemi et al. 2019; Fani and Kohanteb 2017; Palombo and others 2011).

However, the diverse methodologies, solvents, harvest seasons, and other related factors, complicates precise comparisons between study results. In the study by Roozegar et al., which assessed the antimicrobial activity of *Pistacia atlantica* leaf aqueous extract obtained from the mountains of Ilam province, the mean diameter of the inhibition zone against *Streptococcus mutans* was reported as 25 mm using the Disc diffusion method and 22 m using the Embedding sink diffusion method at the highest concentration of extract (100 mg/ml)(Roozegar et al. 2016). Despite variations in the type of extracts, methods for measuring inhibition zones, and sample sources, these results affirm the antibacterial activity of this plant. Conversely, the findings of Mokhtari et al. regarding the aqueous leaf extract and gum of Pistacia atlantica, tested against *Streptococcus mutans*, indicated the absence of a growth inhibition zone around the wells for both the gum and the extract(Mokhtari, Ahrari, and Hosseini 2021). While the presence of alcohol in the extract of our study may potentially influence the diameter of the inhibition zone, it is worth noting that, according to Mokhtari et al., the impact of extracts on bacteria in the well-plate method can be influenced by their diffusion and penetration capabilities into the agar(Mokhtari, Ahrari, and Hosseini 2021). Research on *Lactobacillus rhamnosus* is less extensive compared to *Streptococcus mutans*, complicating the process of drawing conclusions. The results of our study indicate that the hydroalcoholic extract from *Pistacia atlantica* leaf exhibits antibacterial effects against *Lactobacillus rhamnosus* at higher concentrations, albeit less pronounced than its effect on *Streptococcus mutans*. CHX demonstrated superior efficacy against these bacteria. In comparing study results, it is essential to consider various factors such as the season, climatic conditions, region of plant collection, culture medium, extraction methods, and different concentrations of the extract. These factors can significantly influence the antimicrobial activity of the same plant. Therefore, comparing these results and drawing conclusions must be approached with caution. According to current data, CHX demonstrates acceptable antimicrobial efficacy at lower concentrations against oral pathogens. However, various plant extracts may achieve comparable efficacy to CHX at higher concentrations. It is important to note that the potential side effects, including toxicity of these plant extracts should be thoroughly evaluated in in vivo environments. Additionally, the oral environment is complex and differs from laboratory conditions. Therefore, further experiments are necessary to simulate real-life oral conditions and accurately assess the efficacy of the herbal medications.

## Conclusions

The findings of this study revealed that the hydroalcoholic extract derived from *Pistacia atlantica* leaves displayed antibacterial activity against *Streptococcus mutans* and Lactobacillus rhamnosus, comparable to CHX across most concentrations.

